# Diagnosis with Confidence: Deep Learning for Reliable Classification of Squamous Lesions of the Upper Aerodigestive Tract

**DOI:** 10.1101/2022.12.21.521392

**Authors:** Mélanie Lubrano, Yaëlle Bellahsen-Harrar, Sylvain Berlemont, Sarah Atallah, Emmanuelle Vaz, Thomas Walter, Cécile Badoual

## Abstract

**Background:** Diagnosis of head and neck (HN) squamous dysplasias and carcinomas is critical for patient care cure and follow-up. It can be challenging, especially for grading intraepithelial lesions. Despite recent simplification in the last WHO grading system, the inter- and intra-observer variability remains substantial, particularly for non-specialized pathologists, exhibiting the need for new tools to support pathologists.

**Methods:** In this study we investigated the potential of deep learning to assist the pathologist with automatic and reliable classification of HN lesions following the 2022 WHO classification system. We created, for the first time, a large-scale database of histological samples (>2000 slides) intended for developing an automatic diagnostic tool. We developed and trained a weakly supervised model performing classification from whole slide images (WSI). We evaluated our model on both internal and external test sets and we defined and validated a new confidence score to assess the predictions which can be used to identify difficult cases.

**Results:** Our model demonstrated high classification accuracy across all lesion types on both internal and external test sets (respectively average AUC: 0.878 (95% CI:[0.834-0.918]) and 0.886 (95% CI: [0.813-0.947])) and the confidence score allowed for accurate differentiation between reliable and uncertain predictions.

**Conclusions:** Our results demonstrate that the model, associated with confidence measurements, can help in the difficult task of classifying head and neck squamous lesions by limiting variability and detecting ambiguous cases, taking us one step closer to a wider adoption of AI-based assistive tools.

## Introduction

Head and neck squamous cell carcinomas (HNSCC) stand as a substantial global public health threat, occupying the sixth position among the most prevalent forms of cancer (1). The distressing morbidity and mortality statistics associated with HNSCC are primarily due to late-stage diagnoses and challenging treatment protocols (2–4). Early diagnosis of head and neck (HN) lesions is therefore essential to prevent the progression to invasive carcinoma (5). Yet, the categorization of HN dysplasias remains a contentious issue (6,7) due to the varying terminologies, grading methodologies, and low to moderate reproducibility among pathologists (6,8–12) (Table 1).

**Table 1.**
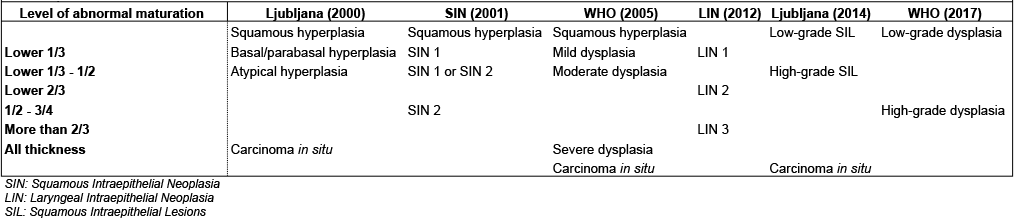
History of dysplasia classification. Overview of the different grading systems for intraepithelial head and neck lesions proposed over the years in the litterature. Table inspired from WHO *Blue Book*

In order to improve inter and intra-rater reliability, the World Health Organization (WHO) recommended in 2017 to grade laryngeal squamous dysplasias with only two categories: low grade and high grade (13,14). This approach indicated that high grade dysplasias carried a ten-fold higher risk of developing into invasive carcinoma compared to low grade lesions (15).

Despite this, reproducibility remains moderate (8) due to the multiple elements to take into account both at cytological and architectural levels, variations in epithelial thickness depending on the anatomical location, and inflammatory and dystrophic changes that often pose challenges in distinguishing true dysplasia (13). Moreover, imposing classification categories on a continuous spectrum of lesions without clear boundaries further exacerbates subjectivity. Given these complexities, the field is in need of new tools to help pathologists make robust and consistent classifications of squamous HN lesions. This would offer physicians better guidance for patients’ monitoring and treatment modalities.

Numerous artificial intelligence (AI) algorithms have been created to support pathologists’ accuracy and consistency (16–25). The classification of HN squamous lesions could significantly benefit from computer-assisted analysis, facilitating standardization and bias reduction in grading. However, research focusing on dysplasia grading, particularly in HN pathology, remains limited. Most studies to date have primarily employed classical machine learning methods rather than deep learning (4,26), concentrating mostly on the oral cavity and not addressing laryngeal lesions (26). The lack of studies in this field could be explained by the absence of public databases including dysplastic lesion annotations, and by the difficulty to achieve grading consensus.

For effective integration into pathologists’ workflows, AI models should provide a confidence estimate for each prediction. While the concept of AI model uncertainty has been studied extensively in the past years (27–29), its application to computational pathology has been sparse (30,31). Tempering the AI model’s predictions with a measure of its confidence could help pathologists to better blend them into their routine grading, thereby fostering model acceptance.

In this study we developed a fully automated weakly supervised model for accurate diagnosis of dysplasias and squamous cell carcinomas of the HN following the current WHO grading system. Figure 1 summarizes the pipeline. The proposed model aims to assist pathologists in achieving automatic and reliable classification of these lesions, with the added benefit of a confidence evaluation of the model’s predictions. We evaluated its performance on a reviewed internal test set as well as an external dataset and proposed a competitive solution to assess the model’s confidence, making it a true tool of peer-review, and thus helping pathologists in their clinical routine.

**Figure 1.**
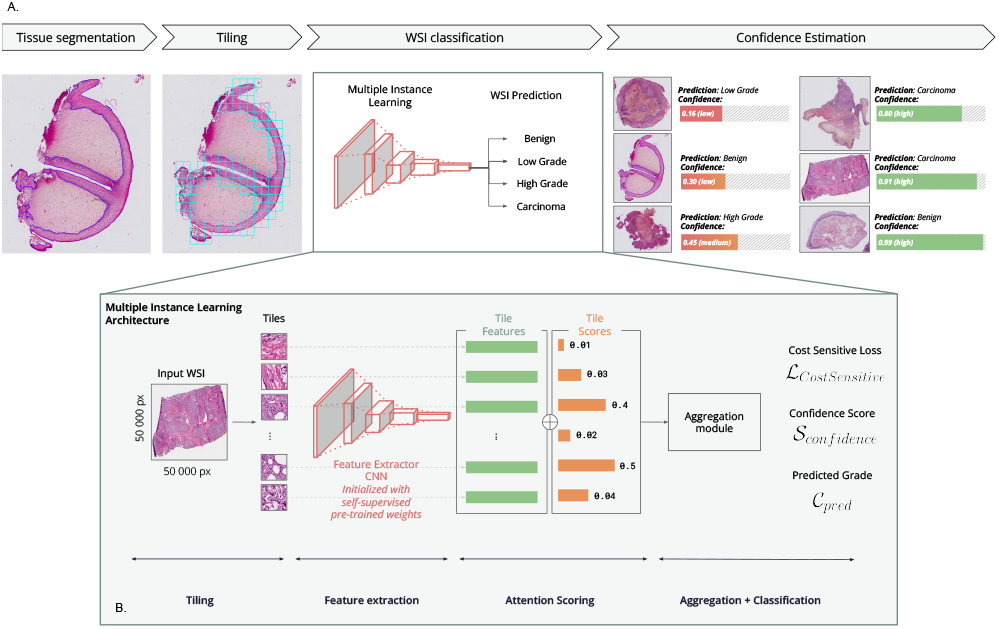
a. Description of the AI-assistive tool pipeline. The AI assistive tool relies on several steps: 1) The tissue of interest (epithelial tissue) is selected with a segmentation model. 2) Small tiles of 224×224 pixels are extracted from the selected areas. 3) Multiple Instance Learning architecture is used to performed WSI classification, choosing between four grades: non-dysplastic, low grade dysplasia, high grade dysplasia or invasive carcinoma. 4) Confidence of the prediction is evaluated thanks to a relevant confidence score. **b. Description of the deep learning model -** 1) Small tiles are extracted from the WSI. 2) The tiles are fed to a frozen feature extractor initialized with pre-trained weights. Relevant features are thus extracted from each tile, resulting in a list of feature vectors of dimension [1, 1024] (the tile’s features). 3) The feature vectors are scored with an attention-based scoring module according to their importance for the downstream classification. 4) An aggregation module performs the weighted sum between the features vectors and the attention scores, and use the resulting vector to perform the final classification. A cost-sensitive loss is used to take into account the ordinal nature of the classes. A confidence score is computed from the outputed risks from the network.

## Materials and Methods

### Dataset Collection

A total of 557 patients were selected retrospectively from 2000 to 2013 from the Hôpital Européen Georges Pompidou (HEGP, Paris, France) clinical database. Patients were at least 18 years old and diagnosed with HN squamous cell dysplasia or carcinoma in the larynx or the pharynx (oropharynx and nasopharynx excluded). Both biopsies and surgical samples were included. In cases of multiple slides per sample, we selected the most representative for the lesion. Pathology reports indicating an inability to discern high grade dysplasia from invasive carcinoma due to tangent inclusion were excluded. The samples were stained using Hematoxylin, Eosin and Saffron (HES) at the time of collection.

Slides were digitized at a 20X magnification (Hamamatsu NanoZoomer® s360), resulting in a pixel resolution of 0.45 µm. We excluded slides with either no surface epithelium or significant artifacts, but retained slides with artifacts that did not compromise clear diagnosis.

Each slide was assigned a global label reflecting the most severe lesion in the sample, adhering to the WHO classification and the clinical report. These initial labels were designated by multiple ENT pathologists from 2000 to 2013. Lesions previously labeled as “mild to moderate” dysplasia were collectively re-evaluated by the pathologist investigators and reclassified according to the latest grading system. The total number of slides was 2064. Table 2 presents patient characteristics and a summary of the cohort.

**Table 2.** Cohort Description.

| Characteristics | Training and Validation sets | Gold Standard Sets (after consensus) | External Set |
| --- | --- | --- | --- |
| **Number of patients** | 456 | 101 | 67 |
| Male | 376 (82.5%) | 79 (78.2%) | 14 (20.9%) |
| Female | 80 (17.5%) | 22 (21.8%) | 53 (79.1%) |
| **Number of samples** | 677 | 115 | 72 |
| Biopsies | 460 (67.9%) | 115 (100%) | 71 (98.6%) |
| Surgical resections | 217 (32.1%) | 0 (0%) | 1 (1.4%) |
| **Anatomical localization** |  |  |  |
| Larynx | 562 | 68 | 77 |
| Pharynx | 55 | 3 | 0 |
| Hypopharynx | 87 | 18 | 10 |
| **Number of slides** | 1949 | 115 | 87 |
| Biospies | 1370 (70.3%) | 115 (100%) | 86 (98.9%) |
| Surgical resections | 579 (29.7%) | 0 (0%) | 1 (1.1%) |
| **Diagnosis (worst lesions on the slide)** |  |  |  |
| Benign (Negative for dysplasia or carcinoma) | 527 (27.0%) | 28 (24.3%) | 21 (24.1%) |
| Low Grade Dysplasia | 205 (10.5%) | 21 (18.3%) | 26 (29.9%) |
| High Grade Dysplasia | 602 (30.9%) | 30 (26.1%) | 17 (19.5%) |
| Invasive Carcinoma | 615 (31.6%) | 36 (31.3%) | 23 (26.4%) |
| *including micro-invasive carcinoma* | 149 (24.2%) | 7 (19.4%) | 0 |
| **Vital Status** |  |  |  |
| Alive | 87 (19.1%) | 19 (18.8%) | - |
| Dead | 140 (30.7%) | 33 (32.7%) | - |
| Lost | 229 (50.2%) | 49 (48.5%) | - |

### Deep Learning Model

Our model builds upon the Attention-based Multiple Instance Learning (MIL) architecture by Ilse et al (32). The WSI is decomposed into small images (called tiles), each of which is mapped to a vector by a Neural Network (NN1). These vectors are then weighted by attention scores (NN2) and summed to build the slide representation, from which the grade is predicted (NN3). NN1 is trained by self-supervised learning (SSL), a powerful technique to learn generic vector representations of images. Of note, SSL allows representation learning independently from the grade prediction task (SimCLR (34)). NN2 and NN3 are trained with a cost-aware classification loss (35) to take advantage of the ordinal nature of the classes. The model was trained to estimate class-specific risk, as opposed to posterior probability, with risks determined by a cost matrix (Supplementary, Table B) that penalizes large errors. For further details, please refer to the Supplementary Materials. All model implementations utilized Python 3.8.0 and Tensorflow 2.5.0.

### Training and Validation

A subset of the dataset was kept aside to create a reference standard test set (115 slides). The remaining slides (1949 slides from 456 patients) were divided into five subsets, four of which were used for training and one for validation (cross-validation (CV) scheme). The subsets were sampled randomly but stratified by patient and grade. Hyperparameters were set to minimize the error measured on the validation folds. Consequently, there were 5 training rounds, with different training (4 folds) and validation sets (1 fold), resulting in five unique models. The outputs of these five models are then averaged to build the final output of the system.

Following best practices, we ensured that there was no patient overlap between training, validation and reference standard test sets.

For performance evaluation, we calculated class-wise classification metrics in a one-vs-all manner. Class-wise scores were averaged to compute overall performance. We evaluated performances by comparing model predictions and labels, with metrics reported alongside 95% confidence intervals (CI) via a bootstrapping method. We further examined predictions through heatmaps of attention scores overlaid on WSIs.

### Reference Standard Test Set

Evaluation of AI algorithms can be problematic if the ground truth is likely to contain errors as well. For this, we have defined a reference set to evaluate our model on the best possible ground truth. To fit with the diagnosis use case, only biopsies were included. The data scientist investigators, not taking part in the grading, were tasked with slide selection, ensuring initial class balance and sample independence. Each slide was selected from either independent patients or from patient slides collected several years apart, with a maximum of two per patient. Slides from the same patient within the same year were excluded resulting in 115 slides selected from 101 patients (Table 2). The reference standard label was determined in two rounds. Two experts independently reviewed the selected slides using the EyeDo© platform, without access to clinical information, initial diagnosis, or the other reviewer’s rating. Afterwards, the two raters met during a consensus meeting to thoroughly discuss the slides on which they disagreed. If the disagreement persisted, the slides were excluded. Initial diagnosis extracted from the patient’s records were used as an additional independent review to measure model performance.

### External Test Set

After the validation of the model on the reference test set, we performed an external validation on slides sourced from another hospital (Hôpital Tenon, APHP, Paris). This external test set included 87 slides from 67 patients (Table 2). The selection criterias were consistent with those of the primary dataset, despite a much more recent timeframe of inclusion (samples from 2016 to 2023). The labeling of the slides followed the most severe lesion directly sourced in the pathology report.

### Confidence Score and Analysis of Misclassified Slides

We calculated a confidence score for each model prediction following the methods described in (36) and in the Supplementary Materials. This score was defined as the difference between the two highest risk outputs from the network, with smaller differences indicating diagnostic uncertainty, and larger differences showing greater confidence. To evaluate the relevance of the confidence score in routine practice, such as screening, we established a threshold to exclude uncertain predictions, optimizing this threshold on the validation set to achieve an overall AUC > 0.9. We assessed the model’s performance on the reference standard test set both before and after the exclusion of uncertain predictions. To understand the model’s limitations, we analyzed attention score heatmaps of slides that were incorrectly, yet confidently, classified.

## Results

### Reference Standard Test Set

The dual blind review yielded independent agreement on 71 slides in the initial round. The remaining 47 slides were subjected to a consensus meeting, resulting in the exclusion of three slides and a consensus decision on the 44 others. Consequently, we used 115 slides from 101 patients as the reference standard test set. Supplementary Materials (Table C) contains additional review details.

### Classification Performance of the Deep Learning Model on the Reference Standard Test Set

On the reference standard test set, the model delivered an average AUC of 0.878 (95% CI: [0.801-0.937]) across the four classes and achieved an AUC > 0.8 for all classes and an AUC > 0.9 for carcinoma detection (ROC AUC displayed in Figure 2). Comparatively, when using initial labels as reference for comparison, the average AUC significantly dropped (AUC=0.817 [0.734-0.888]), confirming the effectiveness of the review in reducing labeling noise and enhancing classification. Comparison of the classification performances of the independent reviewer (initial labels) and the model demonstrate a negligible difference in accuracy (less than 0.02) and confidence intervals that significantly overlap, showing that the model performs similarly to an independent reviewer. Table 3 and Figure 3 summarize classification performance and confusion matrices respectively. Analysis of attention heatmaps verified the model’s capability to focus on significant regions of the slide (Figure 4).

**Table 3.** Classification Performances. For each class, all the metrics are computed in a “one vs rest” manner: slides from the class are considered positives and slides from other classes are considered negatives. The average corresponds to the average over the 4 classes. Confusion matrices are shown in Figure 3. Confidence intervals are computed with bootstrapping (10 000 bootstraps). NPV corresponds to the Negative Predictive Value. PPV corresponds to the Positive Predictive Value.

|  |  | AUC [95% CI] | NPV [95% CI] | PPV (Precision) [95% CI] | Sensitivity [95% CI] | Accuracy [95% CI] | Specificity [95% CI] | AUC (Precision/Recall) [95% CI] |
| --- | --- | --- | --- | --- | --- | --- | --- | --- |
| **AI vs Consensus Labels**<br>**(Reference Standard Set)** | **Average (4 classes)** | **0.878 [0.801-0.937]** | **0.883 [0.811-0.945]** | **0.624 [0.439-0.795]** | **0.621 [0.452-0.788]** | **0.822 [0.748-0.887]** | **0.883 [0.815-0.944]** | **0.708 [0.543-0.844]** |
|  | Normal (0) | 0.916 [0.854-0.962] | 0.911 [0.851-0.966] | 0.68 [0.48-0.864] | 0.68 [0.517-0.862] | 0.861 [0.791-0.922] | 0.911 [0.849-0.965] | 0.78 [0.612-0.903] |
|  | Low Grade (1) | 0.827 [0.735-0.898] | 0.882 [0.802-0.95] | 0.4 [0.219-0.572] | 0.545 [0.323-0.75] | 0.757 [0.67-0.835] | 0.806 [0.725-0.882] | 0.504 [0.29-0.706] |
|  | High Grade (2) | 0.827 [0.731-0.905] | 0.819 [0.736-0.892] | 0.571 [0.333-0.8] | 0.414 [0.24-0.593] | 0.774 [0.696-0.844] | 0.895 [0.829-0.956] | 0.633 [0.441-0.795] |
|  | Carcinoma (3) | 0.942 [0.884-0.982] | 0.921 [0.857-0.973] | 0.846 [0.722-0.946] | 0.846 [0.727-0.946] | 0.896 [0.835-0.948] | 0.921 [0.855-0.974] | 0.914 [0.827-0.973] |
|  | Dysplasia (1+2) | 0.869 [0.8-0.927] | 0.797 [0.697-0.889] | 0.745 [0.619-0.86] | 0.745 [0.615-0.863] | 0.774 [0.696-0.844] | 0.797 [0.692-0.887] | 0.849 [0.754-0.922] |
| **Initial Labels vs Consensus**<br>**Labels**<br>**(Reference Standard Set)** | **Average (4 classes)** | **0.796 [0.707-0.882]** | **0.906 [0.841-0.962]** | **0.682 [0.517-0.839]** | **0.686 [0.519-0.847]** | **0.857 [0.791-0.913]** | **0.907 [0.843-0.962]** | **0.721 [0.587-0.833]** |
|  | Normal (0) | 0.856 [0.764-0.935] | 0.943 [0.889-0.988] | 0.714 [0.542-0.879] | 0.8 [0.625-0.95] | 0.887 [0.826-0.939] | 0.911 [0.848-0.966] | 0.779 [0.646-0.887] |
|  | Low Grade (1) | 0.64 [0.532-0.756] | 0.862 [0.787-0.929] | 0.429 [0.222-0.65] | 0.409 [0.207-0.632] | 0.783 [0.704-0.852] | 0.871 [0.8-0.935] | 0.477 [0.287-0.649] |
|  | High Grade (2) | 0.787 [0.695-0.877] | 0.894 [0.826-0.954] | 0.667 [0.486-0.829] | 0.69 [0.519-0.857] | 0.835 [0.765-0.896] | 0.884 [0.811-0.946] | 0.719 [0.579-0.832] |
|  | Carcinoma (3) | 0.903 [0.839-0.961] | 0.924 [0.861-0.976] | 0.917 [0.816-1] | 0.846 [0.727-0.95] | 0.922 [0.87-0.965] | 0.961 [0.912-1] | 0.908 [0.836-0.963] |
|  | Dysplasia (1+2) | 0.806 [0.731-0.877] | 0.828 [0.73-0.918] | 0.784 [0.667-0.894] | 0.784 [0.667-0.894] | 0.809 [0.739-0.878] | 0.828 [0.732-0.918] | 0.832 [0.75-0.9] |
| **AI vs Initial Labels**<br>**(Reference Standard Set)** | **Average (4 classes)** | **0.817 [0.734-0.888]** | **0.853 [0.777-0.922]** | **0.522 [0.353-0.692]** | **0.523 [0.361-0.686]** | **0.774 [0.698-0.848]** | **0.852 [0.777-0.919]** | **0.6 [0.457-0.733]** |
|  | Normal (0) | 0.91 [0.851-0.962] | 0.889 [0.822-0.954] | 0.72 [0.531-0.886] | 0.643 [0.48-0.826] | 0.852 [0.783-0.922] | 0.92 [0.857-0.967] | 0.76 [0.582-0.901] |
|  | Low Grade (1) | 0.707 [0.595-0.798] | 0.847 [0.767-0.92] | 0.267 [0.12-0.445] | 0.381 [0.176-0.589] | 0.696 [0.617-0.774] | 0.766 [0.675-0.848] | 0.291 [0.161-0.457] |
|  | High Grade (2) | 0.716 [0.615-0.812] | 0.755 [0.667-0.839] | 0.333 [0.12-0.538] | 0.233 [0.088-0.391] | 0.678 [0.591-0.765] | 0.835 [0.759-0.908] | 0.458 [0.292-0.617] |
|  | Carcinoma (3) | 0.932 [0.873-0.978] | 0.921 [0.853-0.974] | 0.769 [0.639-0.9] | 0.833 [0.7-0.939] | 0.87 [0.8-0.93] | 0.886 [0.817-0.951] | 0.892 [0.794-0.959] |
|  | Dysplasia (1+2) | 0.829 [0.748-0.899] | 0.766 [0.656-0.864] | 0.706 [0.566-0.84] | 0.706 [0.571-0.83] | 0.739 [0.652-0.826] | 0.766 [0.667-0.871] | 0.804 [0.693-0.892] |
| **AI vs Initial Labels**<br>**(External Set)** | **Average (4 classes)** | **0.886 [0.813-0.947]** | **0.883 [0.814-0.943]** | **0.655 [0.432-0.863]** | **0.629 [0.464-0.786]** | **0.822 [0.741-0.897]** | **0.878 [0.796-0.946]** | **0.722 [0.543-0.873]** |
|  | Normal (0) | 0.89 [0.815-0.948] | 0.833 [0.747-0.916] | 0.6 [0.333-0.857] | 0.429 [0.217-0.647] | 0.793 [0.701-0.885] | 0.909 [0.836-0.971] | 0.65 [0.427-0.842] |
|  | Low Grade (1) | 0.799 [0.7-0.887] | 0.818 [0.705-0.909] | 0.5 [0.324-0.667] | 0.615 [0.407-0.792] | 0.701 [0.598-0.793] | 0.738 [0.618-0.843] | 0.58 [0.381-0.791] |
|  | High Grade (2) | 0.863 [0.755-0.951] | 0.882 [0.805-0.947] | 0.727 [0.444-1] | 0.471 [0.231-0.706] | 0.862 [0.793-0.931] | 0.957 [0.9-1] | 0.674 [0.423-0.859] |
|  | Carcinoma (3) | 0.994 [0.981-1] | 1 [1-1] | 0.793 [0.625-0.929] | 1 [1-1] | 0.931 [0.874-0.977] | 0.906 [0.83-0.971] | 0.983 [0.94-1] |
|  | Dysplasia (1+2) | 0.831 [0.74-0.908] | 0.727 [0.581-0.854] | 0.721 [0.587-0.848] | 0.721 [0.581-0.842] | 0.724 [0.621-0.816] | 0.727 [0.58-0.854] | 0.82 [0.702-0.911] |

**Figure 2.**
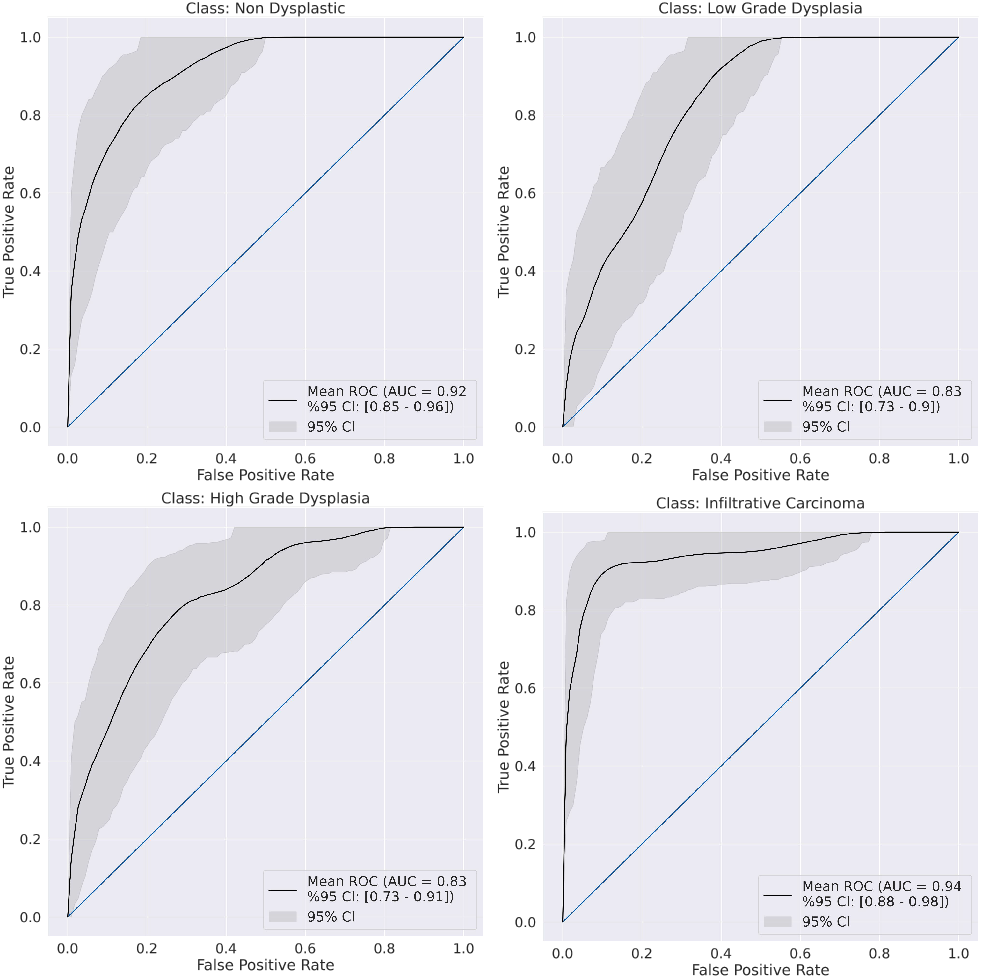
AUC ROC for each class on the reviewed reference standard test set. ROC curves were obtained by bootstrapping of the AI model predictions (10 000 bootstrap samples). They were computed for each class in a One vs Rest manner using consensus labels as a ground truth. ROC = receiver operator characteristic. AUC area under the curve. ROC AUC of the Carcinoma class is better than for the other classes, certainly because the diagnosis of this class is often less ambiguous than for the other grades. Thus, the training data contains less noise on this class, as well as the test data. Misclassification on Carcinoma class concerned microinvasive lesions.

**Figure 3.**
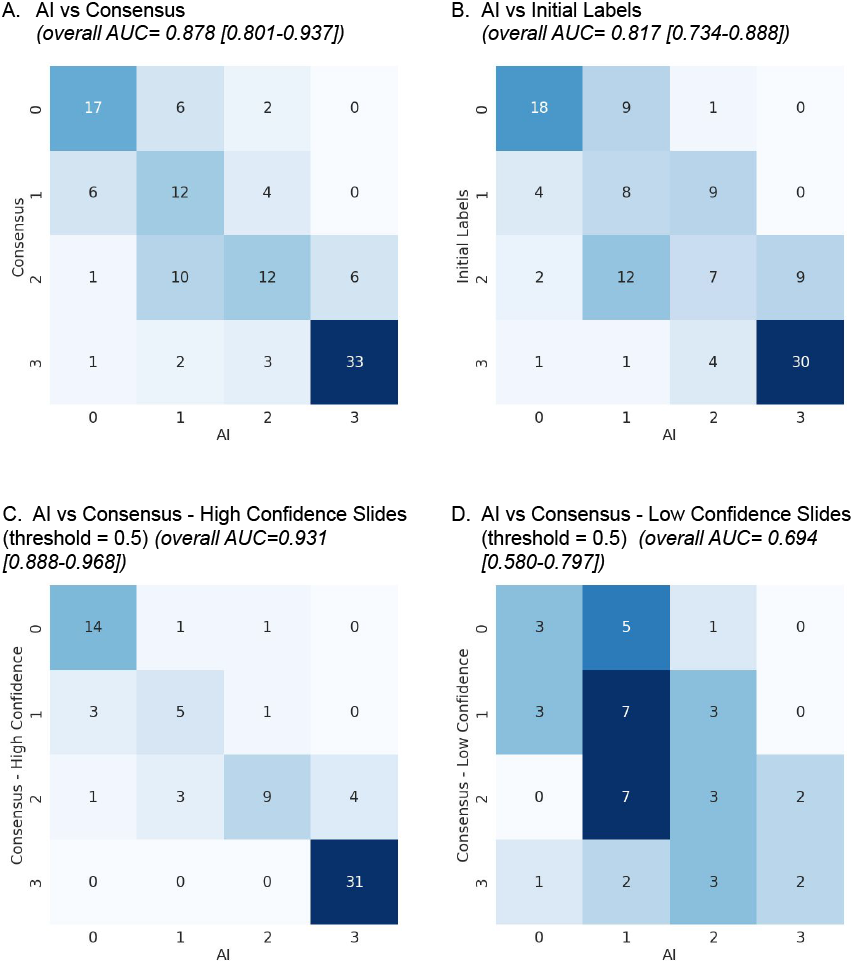
Confusion Matrices. AI model’s performances are evaluated on the reference standard test set on the reviewed labels (A) and the initial labels (from patient’s records) (B). Numbers 0, 1, 2, 3 correspond respectively to classes Non-Dysplastic, Low Grade, High Grade and Carcinoma. Classification performances are superior when using reviewed labels indicating that the review helped reduce noise in the labels. Confusion matrices show that the model is more confused on the Low Grade (1) and High Grade (2) classes, rather than the Carcinoma class (3) for instance which is justified by the ambiguity carried by these classes, on which even pathologists can struggle. Matrix C corresponds to the confusion matrix on the high confident slides at threshold=0.5, matrix D corresponds to the low confident slides. Matrix C is almost diagonal, and the overall AUC on the confident slides subset is higher by more than 10% than on the unconfident subset. Additionally, we see that most of the Carcinoma slides are considered confident by the model. Confusion matrix on the external test set can be found in the Supplementary Materials.

**Figure 4.**
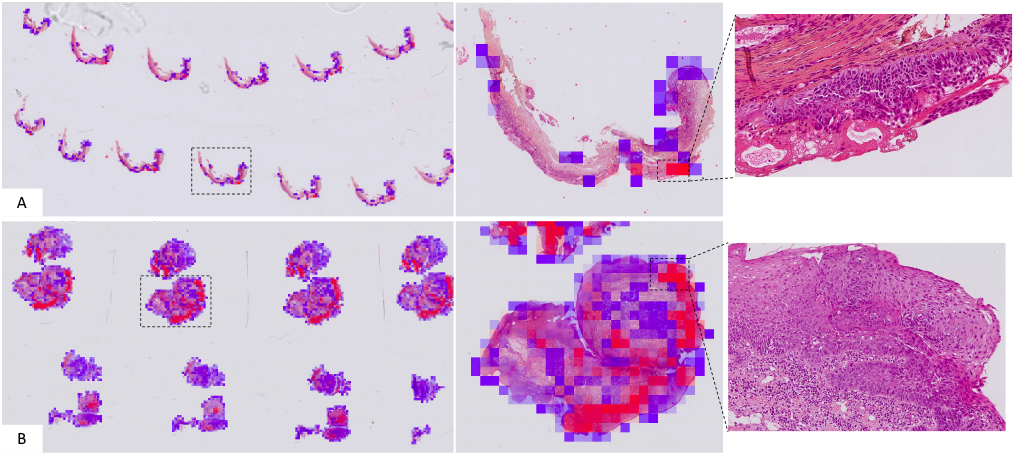
Visualization of Attention Heatmaps. The graphic represents attention scores from the MIL model, overlaid onto WSIs. The color-coding system ranges from 0 (blue) to 1 (red), signifying attention scores, while a lack of color indicates a score of 0.5. The figure showcases the heatmaps of two well predicted slides. The areas with the higher attention scores are displayed on the right under 10X magnification. The slide A (slide_1937) accurately predicted as High Grade Dysplasia, presents a confidence score of 0.43, whereas the slide B (slide_2117), accurately identified as Carcinoma, demonstrates a confidence score of 0.47. The highlighted regions correspond to the regions with marked atypia in the epithelium. Interestingly, heatmaps appeared to be fairly selective, highligting only relevant patterns while displaying low attention scores on normal tissue. Notably, Slide B was discerned by pathologists as containing Micro-Invasive Carcinoma in the initial grading (patient report), and as high grade dysplasia by a reviewer during the dual blind review, marking a challenging diagnosis.

### Confidence Score Assessment

For accurate predictions, the average confidence score was 0.846 +/-0.153, compared to 0.288 +/-0.150 for incorrect predictions. With a 0.5 confidence threshold, 42 slides (36.5%) were marked as uncertain on the reference standard test set, mostly being low grade dysplasias. For the remaining slides, the invasive carcinoma AUC was 0.977 [0.940-1.000], with the model missing no carcinoma slides (Negative Predictive Value of 1.000 [1.000-1.000]). The overall AUC improved by 5.3% (0.931 [0.888-0.968]) when removing slides with low confidence. Conversely, the overall AUC computed on the uncertain slides was equal to 0.694 [0.580-0.797] (−18.4% compared to the overall AUC on the full test set). Figure 3 shows confusion matrices for confident and non confidence slides. Supplementary Figure A suggests the model’s confidence level correlates with the likelihood of reviewer disagreement.

### Analysis of Misclassified Slides

Most misclassifications were low grade dysplasias, labeled as non-dysplastic. Notably, these slides were initially labeled in the clinical record as “non-dysplastic”, showcasing the established ambiguity between these two classes. Four carcinoma slides were misclassified but had low confidence scores, falling beneath the filtering threshold. Three of them had significant artifacts. The other showed carcinoma under a non-dysplastic epithelium, which could be more difficult for the model to identify because of the rarity of this presentation. Misclassified slides associated with high-confidence scores were typically upgraded by one class.

Figure 5 A. shows one high grade dysplasia with a high-confidence score and misclassified as carcinoma. The high-attention tiles displaid severe atypia in the lower epithelium (highlighted by the attention heatmaps). Similarly, Figure 5 B. shows a low grade dysplasia classified as high grade. The two pathologists reviewed the high attention tiles and agreed that these aspects are challenging to interpret.

**Figure 5.**
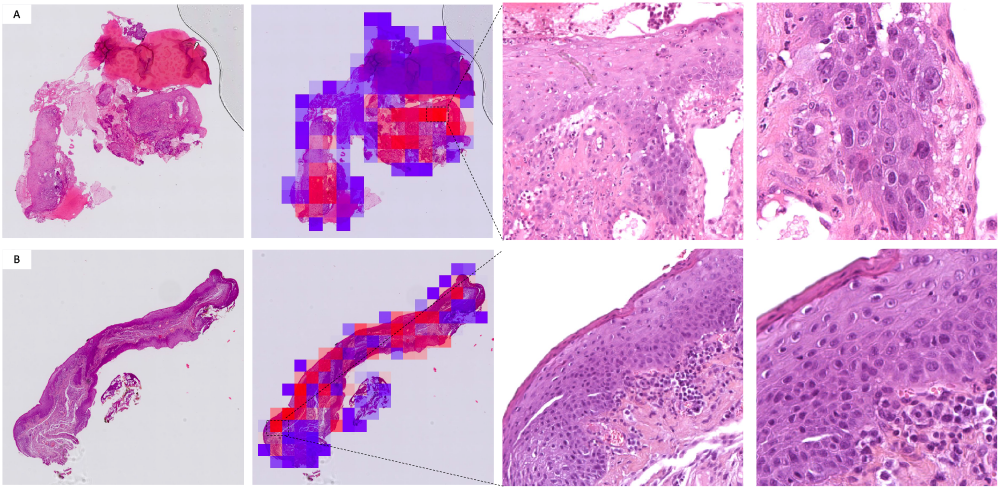
Misclassified slides: attention score analysis. Heatmaps and tiles with high attention scores attributed by the MIL model. First column: one level of a WSI. Second column: WSI overlaid with attention score heatmap. Scores range from 0 (blue) to 1 (red). Third column: 10X magnification. Fourth column: 20X magnification. A.: slide_237. High grade dysplasia predicted as invasive carcinoma. The model focused on marked basal atypia and bulky rete ridges. B.: slide_2712. Low grade dysplasia predicted as high grade. The model focused on marked basal atypia with a corrugated lamina propria. The two lesions are located in the larynx.

### External Validation

On the external validation set the model delivered an average AUC of 0.886 (95% CI: [0.813-0.947]) across the four classes. In particular, the AUC on the Carcinoma class reaches 0.994 (95% CI: [0.981-1]). Similarly as on the internal test set, classification performances reach AUC > 0.8 for all classes, even though the performances are slightly lower on the low grade dysplasia class (class 1) as expected. Confusion matrix is available in the Supplementaries (Figure B).

## Discussion

Our study proposes the first deep learning model for grading HN squamous lesions according to the WHO classification system.

Owing to the absence of a publicly accessible annotated HN dysplasia database, we collected a large scale dataset of HN samples from the HEGP, a prestigious HN diagnosis and healthcare center in France.

We proposed a rigorous reference standard test set for robust model evaluation. The two-phase protocol enabled the creation of a highly reliable ground truth by promoting consensus and encouraging thorough discussion between the two reviewers. By actively engaging in constructive feedback, potential oversight and subjectivity were mitigated. Furthermore, the exclusion of discordant slides yielded a meticulously curated test set. Additionally, we validated our model on an external dataset sourced from a different hospital. This external dataset consisted in slides sampled much more recently (2016-2023) than the primary dataset (2000-2013), providing an additional source of heterogeneity in the data. The labels were sourced from the pathological reports and thus not reviewed. Interestingly, the performances on this external dataset were equivalent if not higher than on the internal reference standard test set. This could be explained by several factors: first, slides being more recent, staining quality was more homogenous and less susceptible to be deteriorated, second, the dysplasias diagnoses (concerning slides from 2017) were made according to the 2017 binary version of the WHO classification, a simpler system that could help diagnosis being more robust and accurate. Finally, the vast heterogeneity and noise in the training data and the use of self-supervised pretraining contributed to make to model robust and widely applicable.

We designed a weakly supervised deep learning model for accurate HN dysplasia grading from annotated WSI, providing an assisted diagnosis tool. Our work introduces a novel confidence score, defined as the difference between the top two class probabilities, ranging from 0 (equally likely) to 1 (maximal confidence). This score was previously validated in a separate study (36), highlighting its relevance in grading precancerous lesions.

This confidence score equips pathologists with a more objective grading method and enhances the practicality and acceptance of AI grading models. High-confidence predictions of severe lesions could be prioritized to reduce delays in diagnosis, while low-confidence cases could be addressed for precautious review. Analysis of false-negative carcinoma samples revealed that technical artifacts causing classification issues also resulted in low confidence scores that would have filtered them out according to our procedure.

In conclusion, our robust deep learning model effectively classifies HN squamous cell non dysplastic epithelium, dysplastic and invasive lesions. Amid a shortage of experts to screen samples or give a second read on difficult interpretations, this model could be used as a powerful and reliable tool by pathologists for faster diagnosis and better grading, ultimately benefiting patient care and follow-up.

## Acknowledgements

Keen Eye team for thoughtful discussions and technical support. Yan Petit for code implementation assistance.

## Author Contributions

Concept and design: CB, TW, SB. Ethical approvals processes : CB, SB. Creation of the clinical and pathology database, slides selection and labeling : YBH. Data management and processing, software implementation, experiments realization and statistical analysis: ML. Choices on experiments : ML and YBH. Design of AI methods: ML, TW. Results analysis: ML and YBH. Discussion of results: ML, YBH, TW, CB. Manuscript writing and review: all authors. All authors read and approved the final paper.

## References

1. Bray F, Ferlay J, Soerjomataram I, Siegel RL, Torre LA, Jemal A. Global cancer statistics 2018: GLOBOCAN estimates of incidence and mortality worldwide for 36 cancers in 185 countries. CA Cancer J Clin. nov 2018;68(6):394–424.

2. Johnson DE, Burtness B, Leemans CR, Lui VWY, Bauman JE, Grandis JR. Head and neck squamous cell carcinoma. Nat Rev Dis Primer. 26 nov 2020;6(1):1–22.

3. Liao LJ, Hsu WL, Lo WC, Cheng PW, Shueng PW, Hsieh CH. Health-related quality of life and utility in head and neck cancer survivors. BMC Cancer. 2019;19(1):1–10.

4. Mahmood H, Shaban M, Indave BI, Santos-Silva AR, Rajpoot N, Khurram SA. Use of artificial intelligence in diagnosis of head and neck precancerous and cancerous lesions: A systematic review. Oral Oncol. nov 2020;110:104885.

5. Mehta N, Tabassum S. Premalignant Conditions of Larynx. In: Pharynx-Diagnosis and Treatment. IntechOpen; 2021.

6. Gale N, Cardesa A, Hernandez-Prera JC, Slootweg PJ, Wenig BM, Zidar N. Laryngeal dysplasia: persisting dilemmas, disagreements and unsolved problems—a short review. Head Neck Pathol. 2020;14(4):1046–51.

7. Hellquist H, Ferlito A, Mäkitie AA, Thompson LDR, Bishop JA, Agaimy A, et al. Developing Classifications of Laryngeal Dysplasia: The Historical Basis. Adv Ther. 1 juin 2020;37(6):2667–77.

8. Mehlum CS, Larsen SR, Kiss K, Groentved AM, Kjaergaard T, Möller S, et al. Laryngeal precursor lesions: Interrater and intrarater reliability of histopathological assessment. The Laryngoscope. oct 2018;128(10):2375–9.

9. Sarioglu S, Cakalagaoglu F, Elagoz S, Ersoy U, Etit D, Hucumenoglu S, et al. Inter-observer Agreement in Laryngeal Pre-neoplastic Lesions. Head Neck Pathol. 21 sept 2010;4(4):276–80.

10. Fleskens SAJHM, Bergshoeff VE, Voogd AC, van Velthuysen MLF, Bot FJ, Speel EJM, et al. Interobserver variability of laryngeal mucosal premalignant lesions: a histopathological evaluation. Mod Pathol. jul 2011;24(7):892–8.

11. Hu Y, Liu H. Diagnostic variability of laryngeal premalignant lesions: histological evaluation and carcinoma transformation. Otolaryngol Neck Surg. 2014;150(3):401–6.

12. Krishnan L, Karpagaselvi K, Kumarswamy J, Sudheendra U, Santosh K, Patil A. Inter-and intra-observer variability in three grading systems for oral epithelial dysplasia. J Oral Maxillofac Pathol JOMFP. 2016;20(2):261.

13. Zidar N, Gale N. Update from the 5th Edition of the World Health Organization Classification of Head and Neck Tumors: Hypopharynx, Larynx, Trachea and Parapharyngeal Space. Head Neck Pathol. 2022;16(1):31–9.

14. El-Naggar AK, Chan JK, Grandis JR, others. WHO classification of head and neck tumours. 2017.

15. Gale N, Blagus R, El-Mofty SK, Helliwell T, Prasad ML, Sandison A, et al. Evaluation of a new grading system for laryngeal squamous intraepithelial lesions—a proposed unified classification. Histopathology. 2014;65(4):456–64.

16. Bera K, Schalper KA, Rimm DL, Velcheti V, Madabhushi A. Artificial intelligence in digital pathology—new tools for diagnosis and precision oncology. Nat Rev Clin Oncol. 2019;16(11):703–15.

17. Bejnordi BE, Veta M, Van Diest PJ, Van Ginneken B, Karssemeijer N, Litjens G, et al. Diagnostic assessment of deep learning algorithms for detection of lymph node metastases in women with breast cancer. Jama. 2017;318(22):2199–210.

18. Steiner DF, MacDonald R, Liu Y, Truszkowski P, Hipp JD, Gammage C, et al. Impact of deep learning assistance on the histopathologic review of lymph nodes for metastatic breast cancer. Am J Surg Pathol. 2018;42(12):1636.

19. Raciti P, Sue J, Ceballos R, Godrich R, Kunz JD, Kapur S, et al. Novel artificial intelligence system increases the detection of prostate cancer in whole slide images of core needle biopsies. Mod Pathol. 2020;33(10):2058–66.

20. Coudray N, Tsirigos A. Deep learning links histology, molecular signatures and prognosis in cancer. Nat Cancer. 2020;1(8):755–7.

21. Bulten W, Litjens G, Pinckaers H, Ström P, Eklund M, Kartasalo K, et al. The PANDA challenge: Prostate cANcer graDe Assessment using the Gleason grading system. 19 mars 2020; Disponible sur: https://zenodo.org/record/3715938

22. Schmauch B, Romagnoni A, Pronier E, Saillard C, Maillé P, Calderaro J, et al. A deep learning model to predict RNA-Seq expression of tumours from whole slide images. Nat Commun. 2020;11(1):1–15.

23. Lu MY, Zhao M, Shady M, Lipkova J, Chen TY, Williamson DF, et al. Deep learning-based computational pathology predicts origins for cancers of unknown primary. ArXiv Prepr ArXiv200613932. 2020;

24. Coudray N, Ocampo PS, Sakellaropoulos T, Narula N, Snuderl M, Fenyö D, et al. Classification and mutation prediction from non–small cell lung cancer histopathology images using deep learning. Nat Med. oct 2018;24(10):1559–67.

25. Courtiol P, Maussion C, Moarii M, Pronier E, Pilcer S, Sefta M, et al. Deep learning-based classification of mesothelioma improves prediction of patient outcome. Nat Med. 2019;25(10):1519–25.

26. Mahmood H, Shaban M, Rajpoot N, Khurram SA. Artificial Intelligence-based methods in head and neck cancer diagnosis: An overview. Br J Cancer. 2021;124(12):1934–40.

27. Gal Y, Ghahramani Z. Dropout as a bayesian approximation: Representing model uncertainty in deep learning. In: international conference on machine learning. PMLR; 2016. p. 1050–9.

28. Lakshminarayanan B, Pritzel A, Blundell C. Simple and scalable predictive uncertainty estimation using deep ensembles. Adv Neural Inf Process Syst. 2017;30.

29. Osband I. Risk versus uncertainty in deep learning: Bayes, bootstrap and the dangers of dropout. In: NIPS workshop on bayesian deep learning. 2016.

30. Pocevičiūtė M, Eilertsen G, Jarkman S, Lundström C. Generalisation effects of predictive uncertainty estimation in deep learning for digital pathology. Sci Rep. 2022;12(1):1–15.

31. Dolezal JM, Srisuwananukorn A, Karpeyev D, Ramesh S, Kochanny S, Cody B, et al. Uncertainty-Informed Deep Learning Models Enable High-Confidence Predictions for Digital Histopathology. ArXiv Prepr ArXiv220404516. 2022;

32. Ilse M, Tomczak J, Welling M. Attention-based deep multiple instance learning. In: International conference on machine learning. PMLR; 2018. p. 2127–36.

33. Huang G, Liu Z, Van Der Maaten L, Weinberger KQ. Densely connected convolutional networks. In: Proceedings of the IEEE conference on computer vision and pattern recognition. 2017. p. 4700–8.

34. Chen T, Kornblith S, Norouzi M, Hinton G. A simple framework for contrastive learning of visual representations. In: International conference on machine learning. PMLR; 2020. p. 1597–607.

35. Chung YA, Lin HT, Yang SW. Cost-Aware Pre-Training for Multiclass Cost-Sensitive Deep Learning. IJCAI. 2016;

36. Lubrano M, Harrar YB, Fick RR, Badoual C, Walter T. Simple and Efficient Confidence Score for Grading Whole Slide Images. In: Medical Imaging with Deep Learning [Internet]. 2023. Disponible sur: https://openreview.net/forum?id=DA1hOTvcMWa

