## Supplementary Materials for "Diagnosis with Confidence: Deep Learning for Reliable Classification of Squamous Lesions of the Upper Aerodigestive Tract"

### Table of Content

**Table of Content**
**Materials and Methods**
Sample inclusion and scanning - Details
Training data management - Details
Deep learning model - Implementation details
Tissue Selection
Binary segmentation model
WSI pre-processing
Description of the Multiple Instance Learning architecture
Table A - MIL Architecture summary
Cost-sensitive training
Table B - Classification Error Costs
Self-Supervised training
Confidence score
**Results**
Reference standard test set
Table C - Consensus meeting results
Confidence score assessment
Figure A - Confidence level distributions
Analysis of misclassified slides
Table D - Misclassified slides
Confusion Matrix on the External Test Set
Figure B - Confusion matrix on the external test set
Comparison between Cost-Sensitive Loss and Cross-Entropy
Table E - Cost Sensitive vs Cross Entropy
**References**

### Materials and Methods

#### Sample inclusion and scanning - Details

The samples were identified in the G Pompidou Hospital's Pathology Department database (Diamic® software) and came from laryngeal and hypopharyngeal locations. Carcinomas sampled after chemotherapy or radiotherapy were excluded since the treatment modifies the aspect of the lesion and the data do not fit the use case. Both biopsies and surgical samples were included, but not all the surgical samples of carcinomas per patient that were available in the hospital's archives. This selection was made in order to keep a balanced proportion of the classes. Every pathology report was read to assess which types of lesions were present. Samples were excluded if the pathologist mentioned in the report that it was impossible for him to distinguish between high-grade dysplasia and invasive carcinoma because of tangent inclusion.

The slides were digitized with a pathology slide scanner (Hamamatsu NanoZoomer® s360) at 20X magnification. All the sections present on the slide were scanned. The quality of the digitization was checked for all the WSIs. Scans were excluded if the digitization failed after two attempts. During digitization, each slide was given an identification number for anonymization purposes.

After digitization, the WSIs were uploaded to the EyeDo© platform. Each slide was given a global label corresponding to the most severe lesion in the sample, following the WHO classification and according to the clinical report. These initial labels were thus provided by several pathologists between 2000 and 2013. Slides with no surface epithelium or with slides with strong artifacts were excluded but slides with artifacts that did not impair a clear diagnosis were kept in the dataset.

#### Training data management - Details

The training data (1949 WSIs from 456 patients) were split into 5 randomly sampled training and validation sets following the 80%-20% standard to train the deep learning model and fix hyperparameters according to best practices. To ensure class balance between training and validation sets, the splits were performed in a stratified manner by patients and grades using the Multilabel Stratified K-Fold algorithm from Sechidis et al (1) (worst grade was kept for each patient). In the training splits, low grades slides were upsampled (1.5 times) to reduce the effect of class imbalance. There was no patient overlap between training, validation and reference standard test sets.

#### Deep learning model - Implementation details

##### Tissue Selection

In order to train the WSI classification model, slides were divided into smaller images of 224x224 pixels at a resolution of 1 µm/pixel. When tiling the entire sample and removing only the white background, WSIs contained up to 8800 tiles (with an average of 1300 tiles per slide), giving a total of 3.9 millions of tiles across the dataset. To reduce the processing time, we removed any tiles not containing epithelial cells (from surface epithelium or tumor). To do so, we first trained a UNet (2) to perform binary segmentation between epithelium and carcinoma tissue (the "foreground") versus any other type of tissue or background (the "background"). The UNet was trained at 10X resolution (1 µm/pixel) on 5439 annotated tiles of 512x512 pixels from 121 slides. The classification threshold was modified to 0.4 in order to minimize false negatives and thus make sure all the tissue of interest was selected. Non-overlapping tiles were then extracted from the selected tissue, resulting in 2.2 millions of tiles.

##### Binary segmentation model

Approximately 600 regions of interest (ROI) of dimensions 4096x4096 pixels were annotated from 121 slides of the dataset. ROI were chosen to maximize the diversity of the tissues. The annotation consisted in detouring epithelial tissues (surface epithelium or carcinoma) resulting in only 1 class (the "foreground"), the rest of the slide was considered as the "background". The annotation was done using the EyeDo© platform. Tiles of 512x512 pixels were extracted from the ROIs at resolution of 1µm/pixel (10X) resulting in 5439 tiles and its associated masks. The tiles were splitted between training, validation and test set respectively to the slides they belonged to. A network derived from UNet (a standard segmentation network architecture) was trained for 100 epochs with early stopping monitoring the validation loss (weighted cross entropy). Data augmentation was used to reduce overfitting. Small detected objects (area < 5000 µm²) were removed from the predictions. Finally, to make sure all the tissues of interest were detected, classification threshold was optimized to reduce the area of false negative tissue and set to 0.4. The segmentation model's performance was assessed through two commonly used metric in image segmentation : the Intersection over Union (IoU) and the Dice Score.

The IoU is computed as the area of overlap between the predicted and actual segmentation divided by the area of union (or combined area) of the predicted and actual segmentation (a higher value indicates a better match between the predicted and actual segmentation). The Dice Score is calculated as twice the area of overlap between the two segmentations divided by the total number of pixels in both images. Both IoU and Dice score ranges from 0 to 1, with 1 indicating perfect agreement between the two segmentations and 0 indicating no overlap. For our model, final Intersection over Union (IoU) on the test set was equal to 0.790 and Dice Score equal to 0.883 reflecting a high level of accuracy in our model's predictions relative to the ground truth.

#### WSI pre-processing

Tiles of 224x224 pixels at resolution of 1 µm/pixel were extracted from the segmented tissue of interest: all the tiles intersecting with foreground detection were kept.

#### Description of the Multiple Instance Learning architecture

Our model was derived from the Attention-based Multiple Instance Learning (MIL) architecture proposed by Ilse et al (3). It consisted of a feature extractor, a scoring module and a classification module. Indeed, due to their size, WSIs have to be cut into smaller tiles. Features are extracted from each tile through a frozen convolutional neural network (DenseNet121, (4)), resulting in feature vectors of dimension 1024. A global label (the grade of the worst lesion on the slide) is associated with the bag of tiles. Thanks to an attention mechanism, the network learns which tiles within the bag are most important for the grading of the lesion and attributes a score. The classification module aggregates the attention scores and the extracted tile representations to obtain a slide representation (weighted sum of the tile representations, where the weights are the attention scores) from which the slide-level label is predicted.

Summary of the attention MIL model can be found in Table A. At each step of the training, one slide with all its tiles was fed to the network. Early stopping was applied and weights were saved at the epochs minimizing the validation loss. ADAM optimizer was used with a learning rate of 1e-4. Dropout layers (rate of 0.7) were inserted between each dense layer and l2 regularization was applied (0.001) to limit overfitting.

#### Table A - MIL Architecture summary

| **Table A -MIL Architecture Summary** | |
|---|---|
| **Layers** | **Type [output dimension]** |
| **Feature Extractor** | DenseNet121 [1024] |
| **Attention-MIL** | **Dimension Reduction**<br>FC [64] + tanh<br>Dropout |
| | **Tile Scoring**<br>FC [64] + softmax<br>Dropout |
| | **Classification**<br>FC [64] + relu<br>Dropout<br>FC [32] + relu<br>Dropout<br>FC [4] |
| *FC: Fully Connected* | |

#### Cost-sensitive training

To leverage the ordinal nature of the class we used the cost aware classification loss introduced in (5): the Smooth One Sided Regression (SOSR) loss.  It allows to train the network to predict class-specific risks rather than posterior probabilities and take into account the order of the classes. The predicted class corresponds to the class minimizing the outputed risk. The SOSR loss is defined as follows:

$$\mathcal{L}_{SOSR} = \sum_i \sum_j ln(1 + exp(\mathbf{2}_{i,j} \cdot (\hat{c}_i - \mathcal{C}_{i,j})))$$

with

$$\mathbf{2}_{i,j} = -\mathbf{1}_{i \neq j} + \mathbf{1}_{i=j}$$

and $\hat{c}_i$ the $i$-th coordinate of the network output and $C$ the error table defined above. The cost matrix used in the loss is described in Table B.

##### Table B - Classification Error Costs

**Table B - Classification Error Costs -** Cost matrix was inspired by the TissueNet Challenge organized by the French Society of Pathology in 2020 that aimed to penalize larger errors than smaller ones. This matrix was designed by a consortium of pathologists prior to the challenge and served as weights for the computation of a custom metric designed for the challenge (weighted accuracy).

| **Table B - Classification Error Costs** | | | | |
|---|---|---|---|---|
| **Ground Truth** | **(0) - Prediction** | **(1) - Prediction** | **(2) - Prediction** | **(3) - Prediction** |
| **Non Dysplastic (0)** | 0.0 | 0.1 | 0.7 | 1.0 |
| **Low Grade dysplasia (1)** | 0.1 | 0.0 | 0.3 | 0.7 |
| **High Grade dysplasia (2)** | 0.7 | 0.3 | 0.0 | 0.3 |
| **Carcinoma (3)** | 1.0 | 0.7 | 0.3 | 0.0 |

##### Self-Supervised training

We initialized the feature extractor with pretrained weights obtained with self-supervised training. SimCLR (6) architecture was used. It consisted of a feature extractor (DenseNet121) and a projection head (3 dense layers). The network was trained on 3.5 million tiles of 336x336 pixels extracted from the entire dataset. The batch size was set to 864, the temperature to 0.1 and the learning rate to 10e-4. To boost the training, the feature extractor was initialized with ImageNet pretrained weights and trained for 5 epochs with frozen feature extractor. After the 5 epochs, the model was trained enterily for 300 hours (143 epochs). Augmentations adapted to histological data were used: crop and resize the tiles to 224x224 pixels, random rotations (90°), flipping, custom stain augmentation and color jittering (brightness, hue, contrast, saturation).

#### Confidence score

Our confidence score is extensively described and benchmarked in (7). We summarize here how it is computed.

It is obtained from the risk estimation vector outputted by the last layer of the network. A softmax is applied to the inverted risk (- risk vector) in order to convert cost estimation into probabilities. The confidence score is then computed by measuring the difference between the two highest probabilities. As the confidence score is derived from the cost sensitive risk estimation, it takes into account the ordinal characteristic of the classes.

The algorithm is described below:

**Algorithm 1 - Confidence score computation**

**Input:** $Y \in \mathbb{R}^n$, with $n$ the number of classes
**Output:** $u$, the confidence score
$Y \leftarrow softmax(-Y)$;
$Y \leftarrow sort(Y, ascending = False)$;
$\mathcal{U} \leftarrow Y_0 - Y_1$;

### Results

#### Reference standard test set

##### Table C - Consensus meeting results

**Table C - Consensus meeting results -** After the 2 reviewers independently reviewed the slides of the reference standard test set, they met for a consensus meeting during which they discussed agreeing on a final grade if there was no agreement in the first place. Slides for which no consensus were found were excluded: one slide for which it was impossible to distinguish high grade dysplasia from invasive carcinoma; one slide for which it was impossible to choose between low grade dysplasia and artifacts; and one slide for which there was a suspicion of carcinoma in the chorion but with no connection to the surface epithelium, which was normal. Five slides (913, 1603, 1627, 2431, 2440, 2591) showed significant discrepancies in grades between the two reviewing pathologists. Most of these discrepancies resulted from reviewer B's tendency to assign lower grades compared to reviewer A and the initial label, although this under-grading didn't occur on carcinoma slides. The slide 454 was initially diagnosed as microinvasive carcinoma, a diagnosis that's particularly challenging due to the lack of consensus on this type of lesions.

| Slide ID | Initial label | Reviewer A (CB) | Reviewer B (YBH) | Consensus | Final label |
|---|---|---|---|---|---|
| 2810 | Dysplasia - high grade | Dysplasia - low grade | Dysplasia - high grade | Yes | Dysplasia - high grade |
| 325 | Non Dysplastic | Dysplasia - low grade | Non Dysplastic | Yes | Dysplasia - low grade |
| 339 | Dysplasia - high grade | Dysplasia - low grade | Non Dysplastic | Yes | Dysplasia - high grade |
| 421 | Dysplasia - high grade | Dysplasia - high grade | Dysplasia - low grade | Yes | Dysplasia - high grade |
| 435 | Dysplasia - high grade | Dysplasia - low grade | Dysplasia - high grade | Yes | Dysplasia - low grade |
| 454 | Carcinoma | Dysplasia - low grade | Dysplasia - high grade | Yes | Dysplasia - low grade |
| 577 | Dysplasia - high grade | Dysplasia - low grade | Dysplasia - high grade | Yes | Dysplasia - high grade |
| 774 | Dysplasia - low grade | Dysplasia - high grade | Dysplasia - low grade | Yes | Dysplasia - high grade |
| 775 | Dysplasia - low grade | Non Dysplastic | Dysplasia - low grade | Yes | Non Dysplastic |
| 776 | Dysplasia - high grade | Dysplasia - low grade | Dysplasia - high grade | Yes | Dysplasia - low grade |
| 126 | Dysplasia - low grade | Carcinoma | Dysplasia - high grade | Yes | Carcinoma |
| 858 | Carcinoma | Non Dysplastic | Dysplasia - high grade | No | Excluded |
| 913 | Non Dysplastic | Non Dysplastic | Dysplasia - high grade | Yes | Dysplasia - low grade |
| 944 | Non Dysplastic | Non Dysplastic | Dysplasia - low grade | Yes | Non Dysplastic |
| 127 | Non Dysplastic | Dysplasia - low grade | Dysplasia - high grade | Yes | Dysplasia - low grade |
| 1279 | Dysplasia - high grade | Carcinoma | Dysplasia - high grade | Yes | Carcinoma |

| | | | | | |
|---|---|---|---|---|---|
| 1282 | Non Dysplastic | Dysplasia - low grade | Non Dysplastic | Yes | Non Dysplastic |
| 1008 | Dysplasia - high grade | Dysplasia - low grade | Non Dysplastic | Yes | Dysplasia - low grade |
| 139 | Dysplasia - low grade | Dysplasia - low grade | Dysplasia - high grade | Yes | Dysplasia - high grade |
| 1403 | Carcinoma | Dysplasia - high grade | Carcinoma | Yes | Dysplasia - high grade |
| 1486 | Dysplasia - low grade | Dysplasia - low grade | Non Dysplastic | Yes | Dysplasia - low grade |
| 1504 | Dysplasia - high grade | Carcinoma | Dysplasia - high grade | Yes | Carcinoma |
| 1594 | Carcinoma | Carcinoma | Dysplasia - high grade | Yes | Carcinoma |
| 1596 | Non Dysplastic | Dysplasia - low grade | Non Dysplastic | Yes | Non Dysplastic |
| 1603 | Dysplasia - high grade | Dysplasia - high grade | Non Dysplastic | Yes | Dysplasia - high grade |
| 1621 | Non Dysplastic | Dysplasia - low grade | Non Dysplastic | Yes | Non Dysplastic |
| 1627 | Dysplasia - high grade | Dysplasia - high grade | Non Dysplastic | Yes | Dysplasia - high grade |
| 1871 | Non Dysplastic | Dysplasia - low grade | Non Dysplastic | Yes | Non Dysplastic |
| 192 | Dysplasia - high grade | Dysplasia - low grade | Dysplasia - high grade | Yes | Dysplasia - low grade |
| 1920 | Non Dysplastic | Dysplasia - low grade | Non Dysplastic | Yes | Dysplasia - low grade |
| 1921 | Non Dysplastic | Dysplasia - low grade | Non Dysplastic | Yes | Non Dysplastic |
| 1971 | Non Dysplastic | Dysplasia - low grade | Non Dysplastic | Yes | Non Dysplastic |
| 2035 | Non Dysplastic | Dysplasia - low grade | Non Dysplastic | Yes | Dysplasia - low grade |
| 2117 | Carcinoma | Carcinoma | Dysplasia - high grade | Yes | Carcinoma |
| 2139 | Non Dysplastic | Dysplasia - low grade | Non Dysplastic | Yes | Dysplasia - low grade |
| 2225 | Dysplasia - low grade | Dysplasia - low grade | Non Dysplastic | Yes | Dysplasia - low grade |
| 2292 | Dysplasia - low grade | Dysplasia - low grade | Non Dysplastic | Yes | Dysplasia - low grade |
| 2294 | Dysplasia - low grade | Dysplasia - low grade | Non Dysplastic | Yes | Dysplasia - low grade |
| 2387 | Dysplasia - high grade | Dysplasia - low grade | Dysplasia - high grade | Yes | Dysplasia - high grade |
| 240 | Non Dysplastic | Dysplasia - low | Non Dysplastic | Yes | Dysplasia - low |

| | | | | | |
|---|---|---|---|---|---|
| | | grade | | | grade |
| 2431 | Dysplasia - high grade | Dysplasia - high grade | Non Dysplastic | Yes | Dysplasia - high grade |
| 2440 | Dysplasia - high grade | Dysplasia - low grade | Non Dysplastic | Yes | Dysplasia - low grade |
| 2442 | Dysplasia - low grade | Dysplasia - low grade | Non Dysplastic | Yes | Dysplasia - low grade |
| 2476 | Dysplasia - high grade | Non Dysplastic | Dysplasia - high grade | Yes | Dysplasia - high grade |
| 2504 | Dysplasia - low grade | Dysplasia - low grade | Dysplasia - high grade | Yes | Dysplasia - high grade |
| 251 | Dysplasia - high grade | Dysplasia - high grade | Carcinoma | Yes | Carcinoma |
| 253 | Dysplasia - high grade | Dysplasia - high grade | Carcinoma | No | Excluded |
| 2591 | Dysplasia - low grade | Dysplasia - high grade | Non Dysplastic | Yes | Dysplasia - low grade |
| 2681 | Dysplasia - low grade | Dysplasia - high grade | Dysplasia - low grade | Yes | Dysplasia - high grade |
| 2682 | Dysplasia - low grade | Dysplasia - low grade | Non Dysplastic | Yes | Dysplasia - low grade |
| 27 | Dysplasia - high grade | Carcinoma | Dysplasia - high grade | Yes | Non Dysplastic |

#### Confidence score assessment

##### Figure A - Confidence level distributions

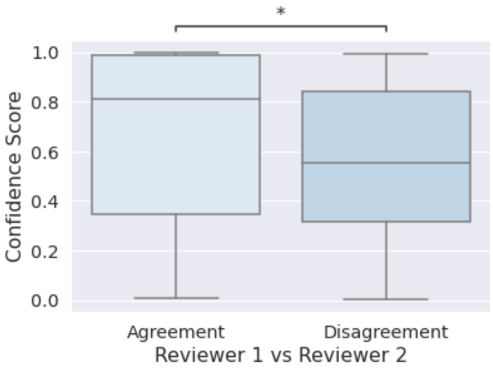


**Figure A - Confidence level distributions -** Comparison of confidence level and agreement between Reviewer 1 and Reviewer 2. The model is more confident on slides on which reviewers agreed during the dual blind review and is less confident on slides on which they disagreed. This suggests that the confidence score reflects the difficulties inherent to the slides, as pathologists would experiment with it. Significance: Mann-Whitney-Wilcoxon test two-sided with Bonferroni correction, p-value=1.095e-02 U-stat=2.455e+03.

#### Analysis of misclassified slides

##### Table D - Misclassified slides

**Table D - Misclassified slides -** Classification probabilities outputted by the network for the misclassified slides of the reference standard test set. The "Filtered-out" column indicates if the slide would have been filtered-out according to our confidence threshold.

**Table C - Misclassified slides**

| Slide ID | Ground Truth | Prediction | Proba class 0 | Proba class 1 | Proba class 2 | Proba class 3 | Confidence Score | Filtered-out |
|---|---|---|---|---|---|---|---|---|
| 126 | 3 | 2 | 0.000 | 0.000 | 0.511 | 0.489 | 0.022 | Yes |
| 856 | 3 | 2 | 0.000 | 0.001 | 0.524 | 0.475 | 0.050 | Yes |
| 1991 | 3 | 0 | 0.528 | 0.470 | 0.002 | 0.000 | 0.058 | Yes |
| 1776 | 0 | 1 | 0.362 | 0.424 | 0.213 | 0.002 | 0.062 | Yes |
| 1008 | 1 | 0 | 0.541 | 0.449 | 0.010 | 0.000 | 0.091 | Yes |
| 2615 | 2 | 3 | 0.001 | 0.007 | 0.450 | 0.542 | 0.092 | Yes |
| 716 | 2 | 3 | 0.000 | 0.001 | 0.443 | 0.555 | 0.112 | Yes |
| 2482 | 2 | 1 | 0.036 | 0.540 | 0.406 | 0.018 | 0.135 | Yes |
| 1627 | 2 | 1 | 0.374 | 0.511 | 0.102 | 0.013 | 0.137 | Yes |
| 454 | 1 | 2 | 0.016 | 0.238 | 0.459 | 0.287 | 0.173 | Yes |
| 2225 | 1 | 2 | 0.001 | 0.015 | 0.579 | 0.405 | 0.173 | Yes |
| 2236 | 0 | 1 | 0.387 | 0.567 | 0.045 | 0.001 | 0.180 | Yes |
| 1339 | 0 | 1 | 0.372 | 0.615 | 0.013 | 0.000 | 0.243 | Yes |
| 777 | 3 | 1 | 0.358 | 0.622 | 0.020 | 0.000 | 0.265 | Yes |
| 577 | 2 | 1 | 0.028 | 0.626 | 0.345 | 0.001 | 0.281 | Yes |
| 325 | 1 | 0 | 0.651 | 0.345 | 0.004 | 0.000 | 0.306 | Yes |
| 421 | 2 | 1 | 0.091 | 0.635 | 0.274 | 0.000 | 0.362 | Yes |
| 2294 | 1 | 0 | 0.694 | 0.305 | 0.001 | 0.000 | 0.390 | Yes |
| 232 | 0 | 1 | 0.052 | 0.671 | 0.276 | 0.002 | 0.395 | Yes |
| 2350 | 3 | 1 | 0.122 | 0.635 | 0.238 | 0.005 | 0.397 | Yes |
| 1279 | 3 | 2 | 0.000 | 0.001 | 0.707 | 0.292 | 0.415 | Yes |
| 715 | 0 | 2 | 0.005 | 0.088 | 0.667 | 0.239 | 0.428 | Yes |
| 2387 | 2 | 1 | 0.273 | 0.716 | 0.012 | 0.000 | 0.443 | Yes |
| 1134 | 0 | 1 | 0.253 | 0.712 | 0.034 | 0.000 | 0.459 | Yes |
| 2712 | 1 | 2 | 0.004 | 0.261 | 0.732 | 0.002 | 0.471 | Yes |
| 339 | 2 | 1 | 0.108 | 0.684 | 0.207 | 0.001 | 0.477 | Yes |
| 788 | 2 | 1 | 0.247 | 0.740 | 0.013 | 0.000 | 0.493 | Yes |
| 1603 | 2 | 1 | 0.096 | 0.661 | 0.143 | 0.100 | 0.518 | No |
| 139 | 2 | 1 | 0.037 | 0.760 | 0.202 | 0.001 | 0.557 | No |
| 2645 | 2 | 3 | 0.001 | 0.004 | 0.208 | 0.788 | 0.579 | No |

| | | | | | | | | |
|---|---|---|---|---|---|---|---|---|
| 785 | 2 | 0 | 0.829 | 0.170 | 0.001 | 0.000 | 0.658 | No |
| 2591 | 1 | 2 | 0.001 | 0.164 | 0.833 | 0.002 | 0.669 | No |
| 2033 | 0 | 1 | 0.161 | 0.835 | 0.004 | 0.000 | 0.673 | No |
| 1434 | 0 | 2 | 0.035 | 0.102 | 0.794 | 0.069 | 0.692 | No |
| 913 | 1 | 0 | 0.881 | 0.119 | 0.000 | 0.000 | 0.762 | No |
| 2811 | 2 | 3 | 0.000 | 0.001 | 0.094 | 0.905 | 0.811 | No |
| 237 | 2 | 3 | 0.001 | 0.005 | 0.082 | 0.912 | 0.829 | No |
| 240 | 1 | 0 | 0.934 | 0.066 | 0.000 | 0.000 | 0.868 | No |
| 455 | 2 | 1 | 0.048 | 0.937 | 0.015 | 0.000 | 0.888 | No |
| 2139 | 1 | 0 | 0.967 | 0.033 | 0.000 | 0.000 | 0.933 | No |
| 2810 | 2 | 3 | 0.000 | 0.000 | 0.010 | 0.990 | 0.979 | No |

#### Confusion Matrix on the External Test Set

Figure B - Confusion matrix on the external test set

**Figure B - Confusion matrix on the external test set -** Predictions on the external sets are obtained similarly than on the reference standard test set: the five cross-validated models are ensembled (the predicted probabilities are averaged) to obtain one unique prediction on the test set.

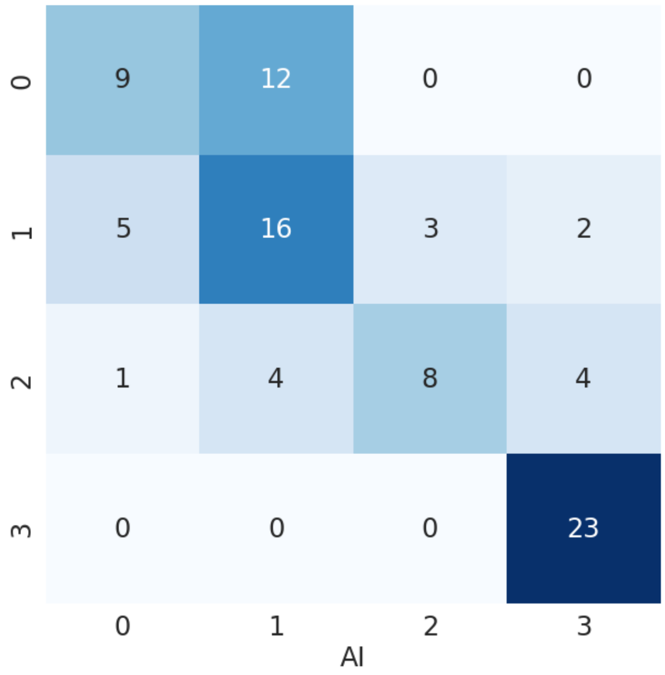

#### Comparison between Cost-Sensitive Loss and Cross-Entropy

##### Table E - Cost Sensitive vs Cross Entropy

**Table E - Comparison between Cost-Sensitive training using the SOSR loss and Cross-Entropy -** For each class, all the metrics are computed in a "one vs rest" manner: slides from the class are considered positives and slides from other classes are considered negatives. The average corresponds to the average over the 4 classes. Confidence intervals are computed with bootstrapping (10 000 bootstraps). We observe a significant gain in term of classification performances when using the cost-sensitive loss. In particular the average AUC gain +6.4% thanks to the cost-sensitive loss. In particular, the cost-sensitive training is more impactful for the intermediate classes, i.e., low-grade and high-grade dysplasia, with respective AUC gains of +14.4% and +10.7%. For the easier classes, namely Non-Dysplastic and Carcinoma, the performance gains are minimal (<1%), indicating that cost-sensitive training plays a significant role in differentiating the intermediate classes.

**Table E - Comparison between Cost-Sensitive training using the SOSR loss and Cross-Entropy**

| | | AUC [95% CI] | NPV [95% CI] | Precision [95% CI] | Recall [95% CI] | Accuracy [95% CI] | Specificity [95% CI] | AUC (Precision/Recall) [95% CI] |
|---|---|---|---|---|---|---|---|---|
| **AI vs Consensus Labels (Cost Sensitive Loss)** | **Average** | **0.878 [0.801-0.937]** | **0.883 [0.811-0.945]** | **0.624 [0.439-0.795]** | **0.621 [0.452-0.788]** | **0.822 [0.748-0.887]** | **0.883 [0.815-0.944]** | **0.708 [0.543-0.844]** |
| | Normal (0) | 0.916 [0.854-0.962] | 0.911 [0.851-0.966] | 0.68 [0.48-0.864] | 0.68 [0.517-0.862] | 0.861 [0.791-0.922] | 0.911 [0.849-0.965] | 0.78 [0.612-0.903] |
| | Low Grade (1) | 0.827 [0.735-0.898] | 0.882 [0.802-0.95] | 0.4 [0.219-0.572] | 0.545 [0.323-0.75] | 0.757 [0.67-0.835] | 0.806 [0.725-0.882] | 0.504 [0.29-0.706] |
| | High Grade (2) | 0.827 [0.731-0.905] | 0.819 [0.736-0.892] | 0.571 [0.333-0.8] | 0.414 [0.24-0.593] | 0.774 [0.696-0.844] | 0.895 [0.829-0.956] | 0.633 [0.441-0.795] |
| | Carcinoma (3) | 0.942 [0.884-0.982] | 0.921 [0.857-0.973] | 0.846 [0.722-0.946] | 0.846 [0.727-0.946] | 0.896 [0.835-0.948] | 0.921 [0.855-0.974] | 0.914 [0.827-0.973] |
| | Dysplasia (1+2) | 0.869 [0.8-0.927] | 0.797 [0.697-0.889] | 0.745 [0.619-0.86] | 0.745 [0.615-0.863] | 0.774 [0.696-0.844] | 0.797 [0.692-0.887] | 0.849 [0.754-0.922] |
| **AI vs Consensus Labels** | **Average** | 0.814 [0.733-0.886] | 0.862 [0.788-0.930] | 0.556 [0.386-0.733] | 0.546 [0.385-0.713] | 0.787 [0.715-0.859] | 0.860 [0.786-0.930] | 0.589 [0.453-0.714] |

| | | | | | | | | |
|---|---|---|---|---|---|---|---|---|
| **(Cross Entropy)** | Normal (0) | 0.913 [0.854-0.963] | 0.890 [0.824-0.954] | 0.750 [0.571-0.920] | 0.643 [0.480-0.826] | 0.861 [0.800-0.922] | 0.931 [0.875-0.977] | 0.758 [0.575-0.901] |
| | Low Grade (1) | 0.683 [0.577-0.779] | 0.847 [0.767-0.920] | 0.267 [0.118-0.455] | 0.381 [0.176-0.589] | 0.696 [0.617-0.774] | 0.766 [0.678-0.855] | 0.252 [0.148-0.376] |
| | High Grade (2) | 0.720 [0.615-0.820] | 0.779 [0.695-0.859] | 0.450 [0.235-0.667] | 0.300 [0.143-0.469] | 0.722 [0.643-0.809] | 0.871 [0.798-0.939] | 0.443 [0.279-0.613] |
| | Carcinoma (3) | 0.941 [0.885-0.983] | 0.932 [0.868-0.986] | 0.756 [0.619-0.889] | 0.861 [0.740-0.971] | 0.870 [0.800-0.930] | 0.873 [0.793-0.949] | 0.902 [0.809-0.967] |
| | Dysplasia (1+2) | 0.829 [0.750-0.900] | 0.769 [0.661-0.868] | 0.720 [0.579-0.848] | 0.706 [0.564-0.836] | 0.748 [0.670-0.826] | 0.781 [0.681-0.885] | 0.800 [0.684-0.892] |

### References


1. Sechidis K, Tsoumakas G, Vlahavas I. On the stratification of multi-label data. In: Joint European Conference on Machine Learning and Knowledge Discovery in Databases. Springer; 2011. p. 145-58.
2. Ronneberger O, Fischer P, Brox T. U-net: Convolutional networks for biomedical image segmentation. In: International Conference on Medical image computing and computer-assisted intervention. Springer; 2015. p. 234-41.
3. Ilse M, Tomczak J, Welling M. Attention-based deep multiple instance learning. In: International conference on machine learning. PMLR; 2018. p. 2127-36.
4. Huang G, Liu Z, Van Der Maaten L, Weinberger KQ. Densely connected convolutional networks. In: Proceedings of the IEEE conference on computer vision and pattern recognition. 2017. p. 4700-8.

5. Y.-A. Chung, H.-T. Lin, and S.-W. Yang. Cost-Aware Pre-Training for Multiclass Cost-Sensitive Deep Learning. IJCAI, 2016.
6. Chen T, Kornblith S, Norouzi M, Hinton G. A Simple Framework for Contrastive Learning of Visual Representations. 13 févr 2020; Disponible sur: https://arxiv.org/abs/2002.05709v3
7. Lubrano M, Bellahsen-Harrar Y, Fick R, Badoual C, Walter T. Simple and Efficient Confidence Score for Grading Whole Slide Images. arXiv preprint arXiv:2303.04604. 2023 Mar 8.
